# APOBEC3A drives metastasis of high-grade serous ovarian cancer by altering epithelial-to-mesenchymal transition

**DOI:** 10.1101/2024.10.25.620297

**Authors:** Jessica M. Devenport, Thi Tran, Brooke R. Harris, Dylan F. Fingerman, Rachel A. DeWeerd, Lojain Elkhidir, Danielle LaVigne, Katherine Fuh, Lulu Sun, Jeffrey J. Bednarski, Ronny Drapkin, Mary Mullen, Abby M. Green

**Affiliations:** Department of Pediatrics, Washington University School of Medicine, Saint Louis, MO; Cancer Biology Graduate Program, Washington University School of Medicine, Saint Louis, MO; Molecular Genetics and Genomics Graduate Program, Washington University School of Medicine, Saint Louis, MO; Division of Gynecologic Oncology, Department of Obstetrics and Gynecology, Siteman Cancer Center, Washington University School of Medicine, St. Louis, MO, USA; Department of Obstetrics, Gynecology, and Reproductive Sciences, University of California San Fransisco, San Fransisco, CA; Division of Anatomic and Molecular Pathology, Washington University School of Medicine, Saint Louis, MO; Penn Ovarian Cancer Research Center, Department of Obstetrics and Gynecology, Perelman School of Medicine, University of Pennsylvania, Philadelphia, PA USA; Basser Center for BRCA, Abramson Cancer Center, Perelman School of Medicine, University of Pennsylvania, Philadelphia, PA USA; Center for Genome Integrity, Siteman Cancer Center, Washington University School of Medicine, St. Louis, MO

## Abstract

High-grade serous ovarian cancer (HGSOC) is the most prevalent and aggressive histological subtype of ovarian cancer, and often presents with metastatic disease. The drivers of metastasis in HGSOC remain enigmatic. APOBEC3A (A3A), an enzyme that generates mutations across various cancers, has been proposed as a mediator of tumor heterogeneity and disease progression. However, the role of A3A in HGSOC has not been explored. Through analysis of genome sequencing from primary HGSOC, we observed an association between high levels of APOBEC3 mutagenesis and poor overall survival. We experimentally addressed this correlation by modeling A3A activity in HGSOC cell lines and mouse models which resulted in increased metastatic behavior of HGSOC cells in culture and distant metastatic spread *in vivo*. A3A activity in both primary and cultured HGSOC cells yielded consistent alterations in expression of epithelial-mesenchymal-transition (EMT) genes resulting in hybrid EMT and mesenchymal signatures, and providing a mechanism for their increased metastatic potential. Our findings define the prevalence of A3A mutagenesis in HGSOC and implicate A3A as a driver of HGSOC metastasis via EMT, underscoring its clinical relevance as a potential prognostic biomarker. Our study lays the groundwork for the development of targeted therapies aimed at mitigating the deleterious impact of A3A-driven EMT in HGSOC.

## INTRODUCTION

Ovarian cancer is the fifth most prevalent cancer among women and is the leading cause of death from gynecologic malignancies worldwide(1). High-grade serous ovarian cancer (HGSOC) represents the most frequent and aggressive histological subtype of ovarian cancer, accounting for approximately 70-80% of ovarian cancer-related deaths(2). HGSOC is often diagnosed at a late stage and exhibits extensive intra- and inter-tumor heterogeneity, resulting in significant challenges in clinical management(3). Intra-tumor heterogeneity in HGSOC manifests as diverse subclones with distinct genomic alterations and gene expression profiles within individual tumors. This clonal diversity is associated with treatment resistance and disease progression(2, 4, 5). Despite aggressive therapeutic interventions, most patients with HGSOC experience relapse and diminished survival, highlighting the critical need for a deeper understanding of the molecular drivers of HGSOC disease progression.

Cancer genome sequencing has established that APOBEC3 (apolipoprotein B mRNA editing enzyme, catalytic polypeptide-like 3) enzymes play a significant role in promoting widespread mutagenesis across various tumor types(4, 6–12). The APOBEC3 family of cytidine deaminases (A3A-A3H) induce mutagenesis through deamination of cytidine to uracil in single-stranded (ss)DNA. Normal function of the APOBEC3 enzymes is as innate immune viral restriction factors(13). Aberrant deaminase activity results in damage to the cellular genome(14–16). Mutations resulting from the enzymatic activity of two APOBEC3 family members, A3A and A3B, leave distinct mutational patterns, defined as single base substitution (SBS) signatures 2 and 13 in the Catalogue of Somatic Mutations in Cancer (COSMIC) database(17). Analysis of tumor genomes has identified APOBEC3 mutagenesis in various cancer types, including breast, bladder, and lung cancer, where is it associated with the accumulation of somatic mutations and the development of subclonal diversity(6, 8, 9, 11, 18–20). Moreover, APOBEC3-mediated mutagenesis has been linked to tumor progression, metastasis, and therapy resistance in breast and lung cancer(19, 21–23).

Detection of APOBEC3 mutational signatures is well established in clear cell ovarian carcinoma(24–28), however prior studies have demonstrated a perplexing range in the contribution of APOBEC3 mutagenesis to overall tumor mutation burden in HGSOC. In a study of more than 100 patients with HGSOC, APOBEC3 mutagenesis was found to contribute to approximately 3% of the overall mutational burden in tumor genomes(28). A smaller study assessing the mutational profiles of various gynecologic cancers found APOBEC3 mutagenesis contributed to a vast range of 0-70% of the overall mutational burden in four patients with HGSOC(25). Thus, the frequency of APOBEC3 mutagenesis in HGSOC remains unclear.

Activity of both A3A and A3B contribute to tumor mutational burden, although several studies have identified A3A as the predominant driver of SBS2 and SBS13 in tumor genomes(8, 29, 30). A germline deletion of A3B, which causes A3A transcript terminating in the A3B 3’UTR, results in increased A3A mutagenesis(31, 32). Epidemiologic studies have demonstrated that this polymorphism is associated with a higher risk of developing tumors, including ovarian cancer(33, 34). The A3B deletion polymorphism is also associated with increased APOBEC3 mutational signatures, indicating elevated A3A activity in tumors(32, 35). However, the biological consequences of A3A activity for HGSOC disease progression are unknown.

In addition to genomic heterogeneity, metastatic progression of HGSOC is driven by epithelial-mesenchymal transition (EMT), a dynamic biological process through which epithelial cells undergo a series of molecular changes that shift the cells towards a mesenchymal phenotype(36). During EMT, cells lose epithelial characteristics, such as cell-cell adhesion and apical-basal polarity, and gain mesenchymal traits, such as motility, invasiveness, and resistance to apoptosis(36). EMT is vital during embryonic development, wound healing, and tissue generation but can be hijacked by cancer cells, facilitating metastasis and disease progression(37, 38). In HGSOC, EMT plasticity generates phenotypically diverse cancer cell populations that exhibit epithelial and mesenchymal characteristics simultaneously, often termed hybrid EMT (hEMT)(39). In a large-scale analysis of EMT phenotypes in cancer, an enrichment of the APOBEC3-mediated SBS2 and SBS13 was observed in tumors exhibiting hEMT expression profiles(38).

In this study, we examine the prevalence of APOBEC3 mutational signatures in HGSOC genomes and investigate the consequences of A3A mutagenesis on patient outcomes. Through genome sequencing of HGSOC, we find an enrichment of A3A mutagenesis in metastastic tumors. Elevated levels of A3A mutagenesis significantly correlates with reduced patient survival. By modeling expression of A3A in HGSOC we demonstrate that A3A activity in HGSOC cells promotes pro-metastatic phenotypes in cultured cells and distant metastatic spread in murine models. Further, we find that A3A activity leads to expression of hybrid EMT and mesenchymal genes in HGSOC cells and metastatic patient tumors, providing a mechanistic explanation for accelerated metastasis. Our study demonstrates the impact of A3A activity on HGSOC progression via EMT, which may be applicable to other tumors.

## RESULTS

### APOBEC3 mutagenesis is enriched in metastatic high-grade serious ovarian cancer

To determine the prevalence of APOBEC3 activity in HGSOC, we assessed genome sequencing from three previously published HGSOC datasets: Pancancer Analysis of Whole Genomes (PCAWG)(40), The Cancer Genome Atlas (TCGA) (41) ^41^, and Dana Farber Cancer Institute/University of Pennsylvania cohort (DFCI/Penn)(42). For each genome, we assessed the contribution of SBS signatures defined by COSMIC. When averaged across each dataset, we found that the APOBEC3 mutational signatures, SBS2 and SBS13, comprised 4.5-6.5% of the mutational burden in HGSOC (**Fig. 1a, Supp. Fig. 1a**).

**Figure 1.**
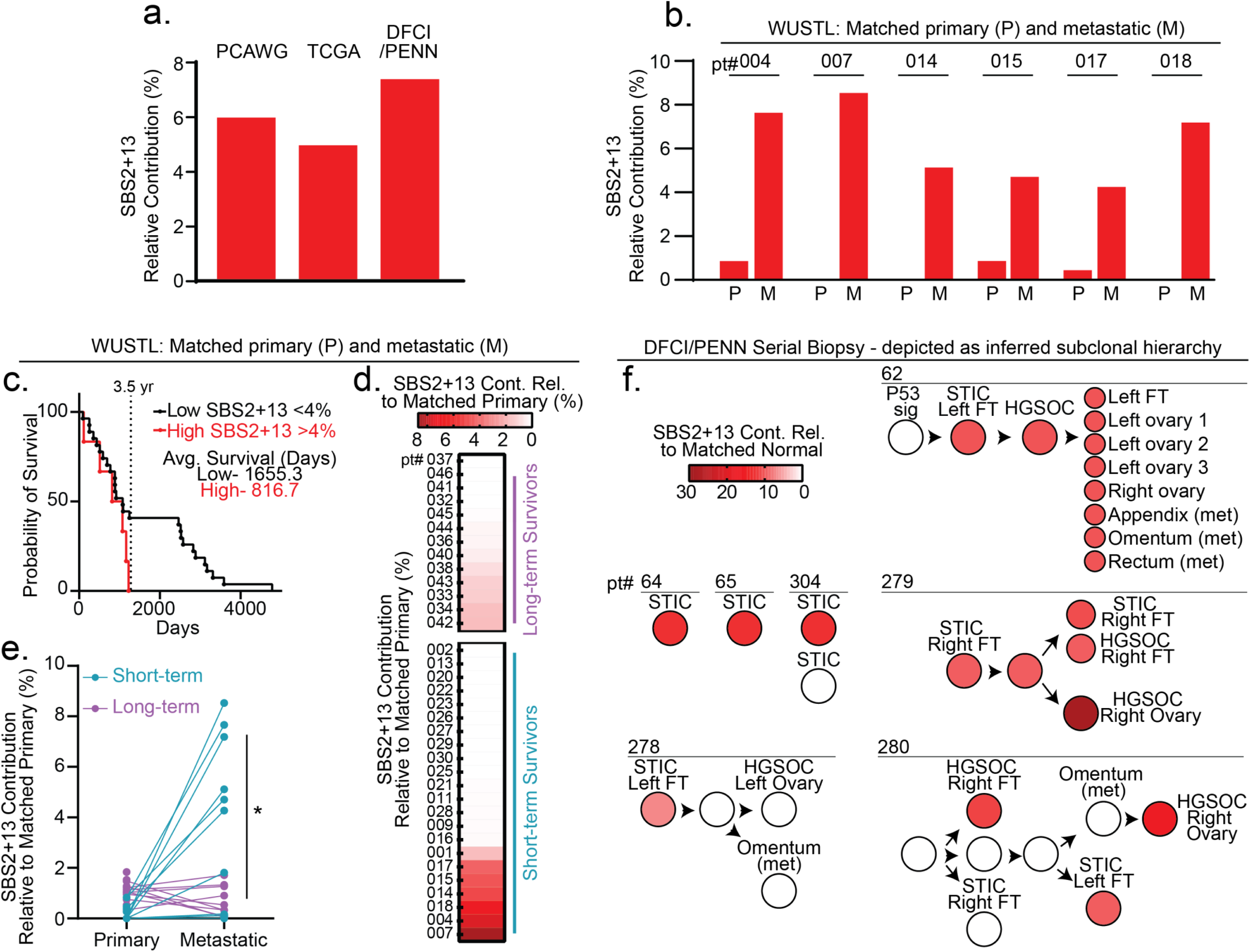
APOBEC3 activity correlates with poor survival in patients with HGSOC. **a**) Whole genome sequencing was assessed to determine the mutational processes occurring within tumor genomes from PCAWG Ovarian Cancer (PCAWG), TCGA Ovarian Cancer (TCGA), and Dana Farber Cancer Institute/University of Pennsylvania (DFCI/PENN) patient cohorts. Relative contribution of APOBEC3 mutational signatures (SBS2 and SBS13) is shown as a fraction of total mutational burden (PCAWG-5.9%, TCGA-4.9%, DFCI/PENN-7.3%). **b-e**) Whole exome sequencing from a patient cohort at Washington University in St. Louis (WUSTL), which includes matched metastatic (M) and primary (P) samples was assessed for relative contribution of SBS2 and SBS13 (**b**). For patients with metastatic sites that demonstrated >4% relative contribution of SBS2 and SBS13 (patient #4, 7, 14, 15, 17, 18), paired P and M SBS2 and SBS13 fractions are shown. **c**) Survival of patients with tumors designated as APOBEC3 high (>4%) vs low (<4%). **d**) Patients were grouped by long-term (>5 years) and short-term (<3.5 years) survival. The relative contribution of SBS2+13 in metastatic sites relative to matched primary is shown for individual patients as a heatmap. **e**) Analysis of relative contribution of SBS2 and SBS13 shows increased SBS2+13 in primary and metastatic sites for individual patients is shown. Blue dots are from patients with short-term survival, purple dots represent patients with long-term survival. **f**) Genomic assessment of multi-site biopsies from patients in the DFCI/PENN cohort. Seven patients with >4% SBS2 and SBS13 relative contribution are shown. Relative contribution of SBS2 and SBS13 shown for each biopsy site ranging from precursor lesions (p53 signatures, serous tubal intraepithelial carcinoma (STIC)) to metastatic sites. Three patients had biopsies of only STIC lesions (64, 65, 304). Unlabeled circles and arrows between lesions indicate inferred subclonal hierarchy, adapted from Labidi-Galy, et al.(42), and predicted progression of disease.

In a previous pan-cancer study, APOBEC3 mutagenesis was determined not to be significantly different between primary tumors and metastatic sites(28). To query whether differences in APOBEC3 activity exist between primary and metastatic sites in HGSOC, we took advantage of a dataset from Washington University in Saint Louis (WUSTL) which includes paired primary and metastatic whole exome sequencing (WES) from 35 patients with HGSOC(43). When pooling all primary and metastatic samples, we observed SBS2 and SBS13 were increased in metastatic sites by 6-fold relative to primary sites (**Supp. Fig. 1b**). We then looked at the individual patients from the WUSTL cohort with the highest contribution of APOBEC3 signature mutations to tumor mutational burden. Six of 35 patients (5.8%) exhibited >4% contribution of APOBEC3 signature mutations in metastatic tumor genomes (**Fig. 1b, Supp. Fig. 1c**). All six patients had an increase in the contribution of APOBEC3 signature mutations in metastatic sites compared to primary tumors (**Fig. 1b**).

Given the observed association between APOBEC3 mutagenesis and tumor metastasis, we investigated how APOBEC3 SBS signatures correlated with patient survival. We divided the WUSTL patients by those with high (>4%) or low (<4%) contribution of APOBEC3 signature mutations in metastatic tumors. We found that patients with high APOBEC3 mutagenesis had an average survival of 816.7 days while patients with low APOBEC3 mutagenesis had an average survival of 1655.3 days, correlating a high burden of APOBEC3 SBS signatures with decreased overall survival (**Fig. 1c**). We next analyzed patients in the WUSTL cohort by categorizing them as short-term (<3.5 years) or long-term (>5 years) survivors(43). We identified that APOBEC3 SBS signatures were more abundant in tumor genomes from short-term compared to long-term survivors (**Fig. 1d**). Upon examination of primary-metastatic pairs, we found that APOBEC3 SBS signatures were enriched in the metastatic sites of short-term survivors compared to long-term survivors (**Fig. 1e**). These data indicate that APOBEC3 mutagenesis is enriched in metastatic HGSOC and correlated with decreased patient survival.

As APOBEC mutagenesis increased from primary to metastatic tumors, we sought to determine the kinetics of APOBEC3 activity throughout HGSOC evolution. Within the DFCI/Penn cohort, we assessed multi-site biopsies from patients ranging from normal fallopian tube tissue, *TP53* mutant single-cell epithelial layer (p53 signature), serous tubal intraepithelial carcinoma (STIC), and primary and metastatic HGSOC lesions(42). In addition, three patients with only STIC lesions were analyzed for mutational signatures relative to normal fallopian tube tissue. Of nine patients in the cohort, seven had measurable APOBEC3 mutational signatures in at least one tumor site (shown in **Fig. 1f**). Interestingly, all seven patients in whom APOBEC3 mutational signatures were evident had measurable APOBEC3 mutational signatures early in tumor development at the STIC lesion stage, with contributions ranging from 7-21%. Four of the DFCI/Penn patients had sequenced biopsies of samples beyond the STIC lesion stage, and 3/4 exhibited persistence or increase of APOBEC3 mutagenesis as tumors evolved (9.5-29% contribution, **Fig. 1f**), consistent with the increased APOBEC3 activity observed in the WUSTL cohort metastatic lesions. These data indicate that APOBEC3 mutagenesis may arise early in HGSOC development but can accelerate or accumulate throughout tumor evolution. Together, findings from primary tumor genomes demonstrate relatively frequent A3A mutagenesis in HGSOC that is enriched in metastatic sites and associated with poor survival suggesting that A3A activity enables tumor progression.

### Modeling episodic A3A expression in HGSOC cells

To experimentally determine how APOBEC3 mutagenesis impacts ovarian cancer progression, we developed cellular models of APOBEC3 expression in HGSOC. We utilized two HGSOC cell lines to model episodic A3A activity. A3A is an interferon (IFN)-stimulated gene(44) and we selected OVCAR3 and OVCAR4 which do not express A3A even when stimulated with IFN (**Supp. Fig. 2**). Importantly, both cell lines have *TP53* inactivating mutations, consistent with the designation of HGSOC. We introduced a doxycycline (dox)-inducible A3A transgene by lentiviral integration into both cell lines (OVCAR3-A3A and OVCAR4-A3A), enabling controllable A3A expression and activity (**Fig. 2a, Supp Fig 3**). A3A mutagenesis has been shown to occur in intermittent bursts over time(45, 46). To mimic intermittent A3A expression in cancer, we treated cells once weekly with low dose (0.5 g/mL) dox followed by dox washout 72 hours later (**Fig. 2a**). This approach resulted in A3A expression after one day of treatment that gradually led to undetectable levels by day five of treatment (**Fig. 2a**). We repeated this treatment course every 7 days for 8 weeks after which we initiated experimental investigations. This treatment schema provided a system to address how historic, intermittent A3A expression, rather than the consequences of ongoing A3A mutagenesis, impacts tumor cell phenotype. To account for the previously established stochastic nature of A3A activity(7), we generated three independent replicates of each cell line (A3A V1, A3A V2, A3A V3) (**Fig. 2b**). All A3A replicates were derived from non-treated (NT) control cells which underwent viral integration of the transgene but were never exposed to dox or A3A expression (**Fig. 2b**).

**Figure 2.**
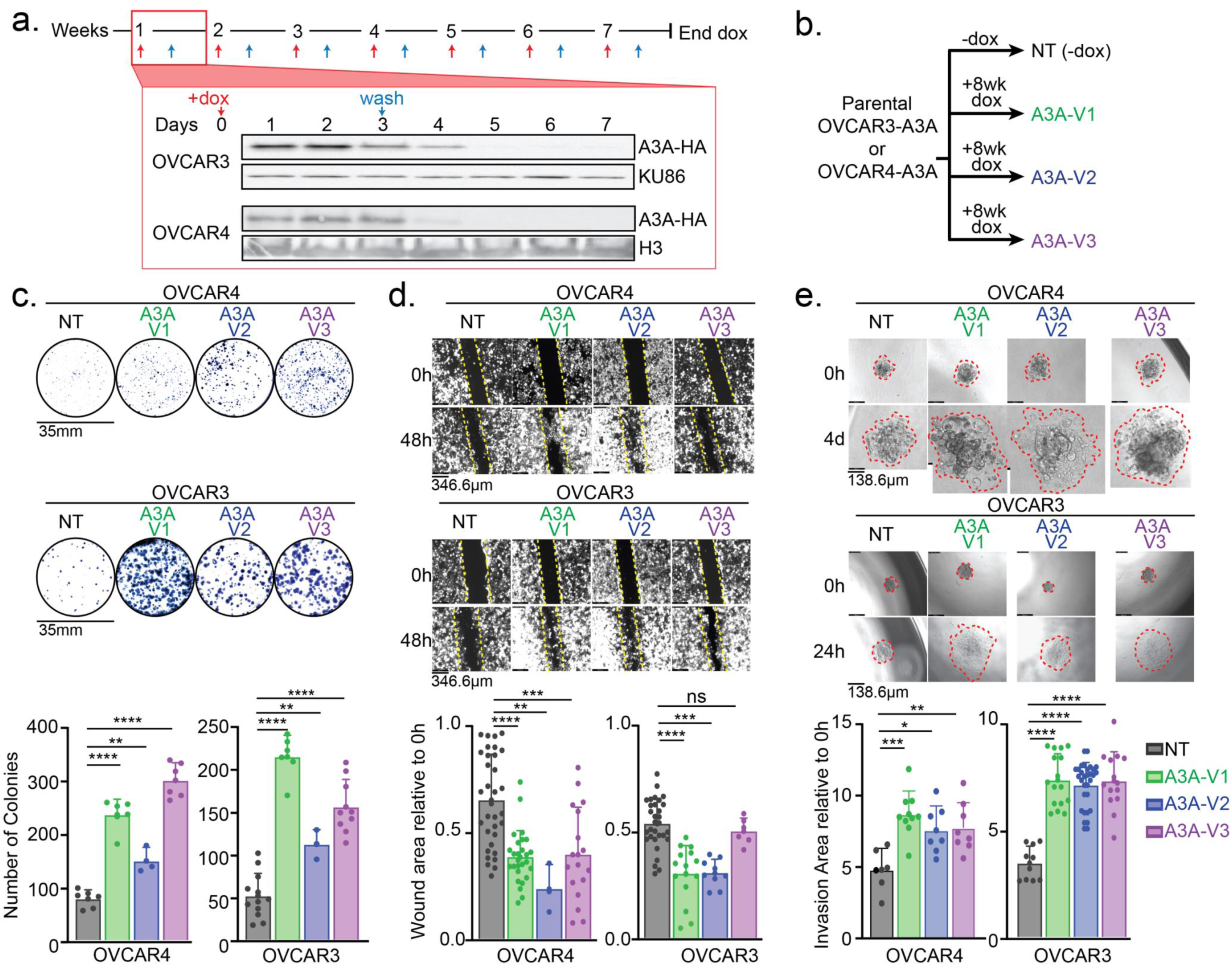
Episodic A3A expression promotes HGSOC cell survival, migration, and invasion. **a**) OVCAR3 and OVCAR4 cells engineered to stably express a dox-inducible A3A transgene were treated with dox on day 0 and washed out on day 3. Treatment schema was repeated for 8 weeks. Immunoblot of HA-tagged A3A from cell lysates harvested on sequential days throughout one week following dox treatment. Ku86 and H3 are loading controls. **b**) Three biological replicates of A3A cell lines (A3A V1-3) were independently derived from the parental cell lines (NT). NT cells were culture in parallel for 8 weeks. **c**) Cell survival under stress was assessed using colony formation assays. Cells were seeded at ultra-low cell densities and resulting colonies were then stained, imaged, and quantified. **d**) Wound healing assays were performed to assess the migratory phenotype of OVCAR3 and OVCAR4 NT and A3A V1-3 cells. The wound was imaged using a 4x objective at 0h and 48h. Wound area at 48h relative to 0h is shown in bar graph. **e**) Spheroids of each cell line were generated and placed onto a Matrigel-containing pseudo-basement membrane. Spheroids were imaged with a 10x objective at 0h, 24h, and 4 days. Area of the spheroid at 24h or 4 days relative to 0h is shown. Invasion area is outlined in red. For data in **c-e** representative images are shown and quantification is depicted in bar graphs below. Significance was determined by unpaired two-tailed t-test, ****p≤0.0001, ***p≤0.001, **p≤0.01, *p≤0.05, ns p>0.05. Error bars are mean with SD for n≥3 replicate experiments. Representative images are shown.

**Figure 3.**
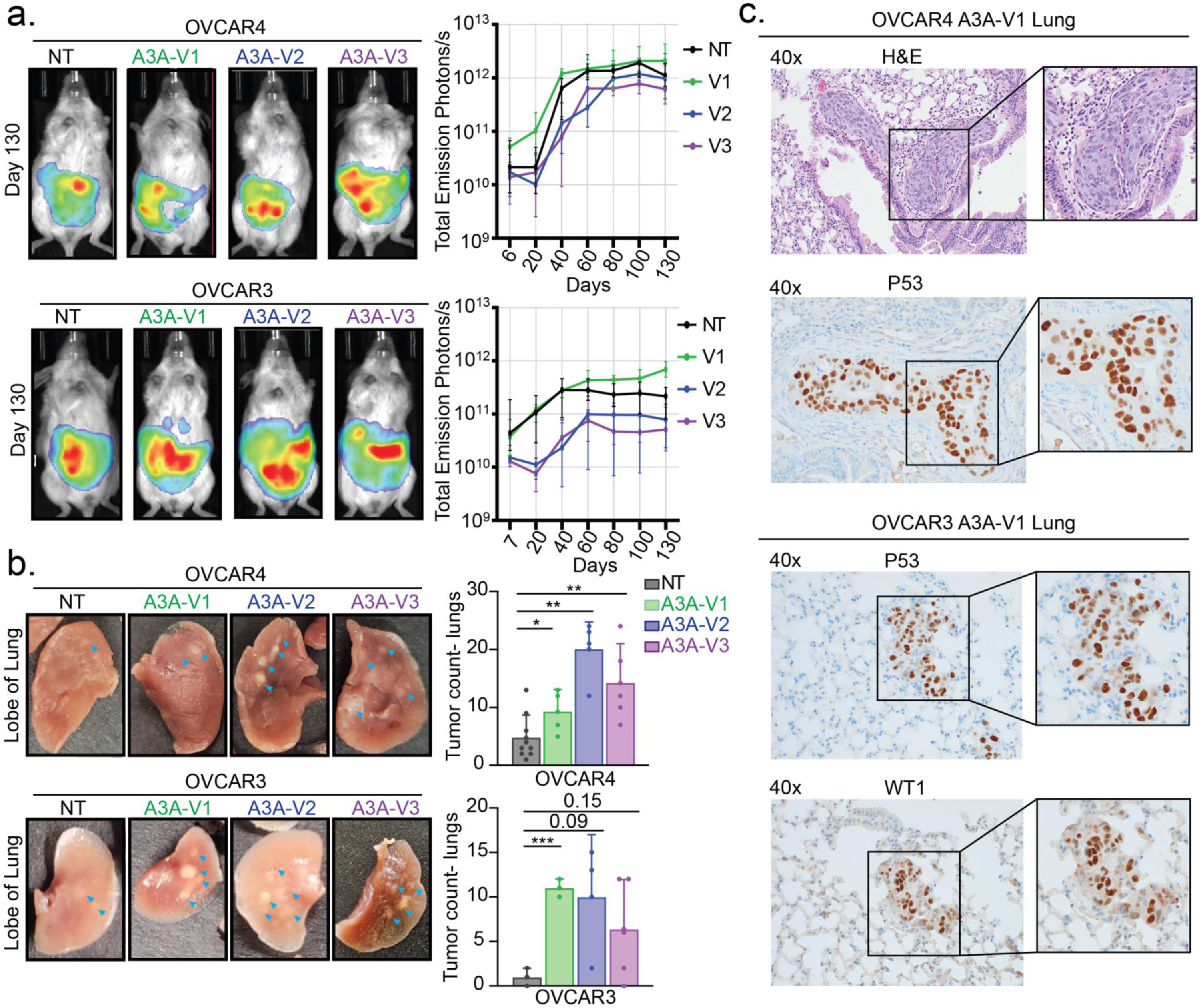
A3A promotes distant HGSOC metastasis *in vivo*. **a**) Representative images of total emission in photons/sec of BLI from day 130 post-injection. BLI over time plotted as total emission in photons/sec for each time point, median and error of replicates are shown. **b**) Lungs were harvested from the mice at death or 165 days and tumor burden was determined by macroscopic count of tumor nodules. Representative images are shown, blue arrows indicate macroscopic tumor sites. Significance determined by two-tailed t-test. Error bars are mean with SD for duplicate experiments (n=3-5 mice for each experiment). ***p≤0.001, **p≤0.01, *p≤0.05. **c**) Lungs were sectioned and analyzed by hematoxylin and eosin (H&E) staining, and immunohistochemical staining for TP53 and Wilms’ tumor-1 (WT1). Representative images are shown at 40x.

### A3A expression promotes HGSOC cell survival, migration, and invasion

As APOBEC3 mutagenesis in patient tumors was correlated with decreased survival (**Fig. 1c-e**), we hypothesized that APOBEC3A would promote phenotypic changes consistent with increased metastatic potential. We examined three key steps in metastasis - cell survival, migration, and invasion - in A3A-exposed cells. We assessed the ability of HGSOC cells to survive under stress by seeding at ultra-low densities. We found that OVCAR3-A3A and OVCAR4-A3A cells formed significantly more colonies than control (NT) cells (**Fig. 2c**). Increased colony formation was not due to an altered rate of proliferation, as A3A and NT HGSOC cells grew at similar rates (**Supp. Fig. 4**) In addition, using wound healing assays we found that OVCAR3-A3A and OVCAR4-A3A cells migrated more rapidly into a wound than NT controls. Cells were grown to confluency, then a scratch wound was created in the plate. Cell migration into the wound was monitored by microscopy. Forty-eight hours after wound formation, both NT and A3A-exposed cells migrated into the wound but A3A-exposed HGSOC cells re-filled a larger area of the defect, consistent with enhanced migration (**Fig. 2d**).

Finally, we assessed how episodic A3A impacts the invasive potential of HGSOC cells. We generated spheroids of each cell line and placed them in a pseudo-basement membrane (50% Matrigel). By serial imaging, we found that A3A-exposed OVCAR3 and OVCAR4 cells demonstrated a greater area of invasion into a pseudo-basement membrane than NT controls (**Fig. 2e**). Notably, we found that all OVCAR3-A3A and OVCAR4-A3A cell lines (V1-V3) exhibited similarly altered phenotypes. Together these data show that episodic A3A expression promotes pro-tumor, metastatic phenotypes in culture indicated by increased survival, migration, and invasion.

### A3A accelerates distant HGSOC metastasis *in vivo*

We reasoned that the phenotypic changes observed in HGSOC cells in culture would impact tumor metastases *in vivo*. As ovarian cancer progresses in patient tumors, clusters of cancer cells detach from the primary tumor into the peritoneal cavity and seed both local and distant sites(47). Thus, we designed *in vivo* experiments to mimic ovarian cancer progression in patients by engrafting OVCAR3 or OVCAR4 cell clusters suspended in 50% Matrigel into the peritoneal cavity of immunodeficient mice. This protocol enabled the formation of tumor spheroids immediately after injection, replicating the behavior of tumor clusters that have migrated from the primary tumor site in patients(47, 48). After delivery of the OVCAR3 or OVCAR4 cells, we monitored tumor burden via bioluminescent imaging (BLI) and found stable engraftment of all cell lines (NT and A3A V1-3) within 2-3 weeks of injection (**Fig. 3a**). Following stable engraftment of tumor in the peritoneal cavity, we monitored tumor burden through serial BLI for 165 days and found no differences in overall tumor burden between A3A-exposed and NT control xenografts (**Fig. 3a**). Both BLI and post-mortem measurement of tumor weights demonstrated a high burden of disease within the peritoneal cavity, consistent with multifocal seeding from intraperitoneal tumor injection. (**Fig. 3a, Supp. Fig 5**).

We next examined extra-peritoneal organs to evaluate distant metastasis. Through macroscopic assessment of tumor nodules in the lungs, we found that mice engrafted with A3A-exposed OVCAR3 and OVCAR4 cells had significantly more metastatic seeding in the lungs than NT controls (**Fig. 3b**). Histopathologic examination and immunohistochemical staining for WT1 and p53 confirmed the presence of ovarian tumor within the lungs(49) (**Fig. 3c**). These data demonstrate that episodic A3A activity in HGSOC cells enhances tumor metastasis to distant sites.

### Episodic A3A activity in HGSOC cells leads to stochastic mutagenesis

Given the distinct pro-metastatic phenotype of A3A-exposed HGSOC cells in culture and *in vivo*, we sought to identify mechanisms underlying A3A-mediated tumor behavior. First, to determine how A3A activity impacts the genomes of OVCAR3 and OVCAR4 cells, we performed deep WES (∼200x) on NT and A3A V1-3. Using NT cells as reference genomes, we assessed how the mutational burden was altered by episodic A3A expression. We found that SBS2 and SBS13 signatures contributed to ∼20-70% of the novel mutational burden after episodic A3A exposure (**Fig. 4a-b**). While the fraction of SBS2 and SBS13 contributing to the overall mutational burden in these cells is higher than we report for patient tumors (**Fig. 1, Supp. Fig. 1**), the total number of mutations caused by A3A (range 81-759) are analogous to those reported in patient tumor sequencing (average of 151 mutations/tumor attributable to SBS2 and SBS13 in ovarian cancers from COSMIC(**Fig. 4c-d**)(17)).

**Figure 4.**
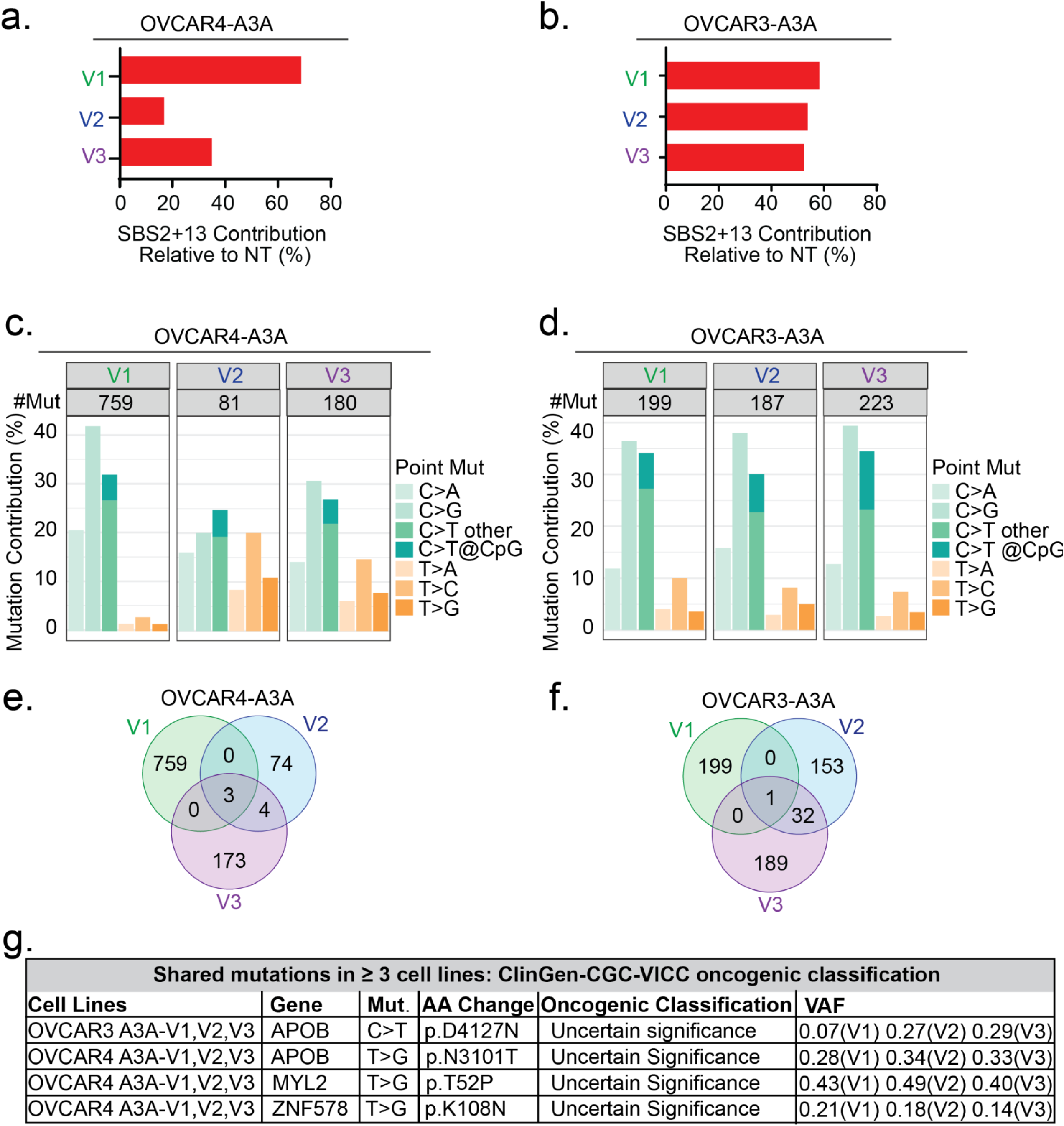
Episodic A3A in HGSOC causes stochastic mutagenesis. Whole exome sequencing of OVCAR4 and OVCAR3 A3A V1-3 and NT cell lines was performed. Respective NT cell lines were used as a reference genome to determine *de novo* mutations in A3A-exposed cells. **a-b**) Contribution of SBS2 and SBS13 to total *de novo* mutations. **c-d)** The total number of *de novo* base substitution mutations and relative contribution of transition and transversion mutations. Total number of acquired mutations is shown for each version. **e-f)** Venn diagram of mutations acquired in A3A-exposed OVCAR4 and OVCAR3 cell lines versions 1-3. **g)** ClinGen-CGC-VICC was used to identify the oncogenic classification of mutations acquired in ≥3 A3A-exposed cell lines. The cell line, mutated gene, base substitution, amino acid change, oncogenic classification, and variant allele frequency (VAF) are shown.

Next, we assessed the novel mutations that were acquired in A3A-exposed cells. We found variation in the number of mutations across versions (**Fig. 4c-d**). The mutations acquired were largely unique to each version (**Fig. 4e-f**, **Supp. Table 1**), consistent with prior findings that A3A acts stochastically on the genome(7, 30, 45). In OVCAR4-A3A V1-3, only 7 mutations were shared between at least two versions and none were classified as likely to be oncogenic by ClinGen-CGC-VICC(50) (**Fig. 4e, g, Supp. Table 1**). In OVCAR3-A3A V1-3 we found 33 mutations shared among at least two versions, none of which were classified as oncogenic (**Fig. 4f-g, Supp. Table 1**). These data suggest that A3A-induced mutations in protein-coding regions did not explain the shared pro-metastatic phenotypes observed in A3A-exposed cells.

### Episodic A3A expression alters the EMT trajectory of HGSOC

Given the lack of shared aberrations in the protein-coding genome to explain the A3A-mediated metastatic phenotype, we sought to assess transcriptional changes that may provide a mechanism for the consistent phenotypic changes observed across A3A-exposed HGSOC cells. From RNA-sequencing, we identified differentially expressed genes (DEGs) in A3A-exposed cells relative to NT controls (**Supp. Fig. 6a-b**). We additionally assessed DEGs from metastatic patient tumors in the WUSTL dataset by comparing those with high versus low burdens of APOBEC3 mutagenesis as defined in **Fig. 1c** (**Supp. Fig. 6c**). Through gene set enrichment analysis, we determined that DEGs in all A3A cell versions were significantly enriched for genes associated with EMT (**Fig. 5a, Supp. Tables 2-3**). We found a similar enrichment of EMT genes in metastatic tumors from patients in the WUSTL dataset that had APOBEC3 high relative to low mutational burden (**Fig. 5b, Supp. Tables 2-3**).

**Figure 5.**
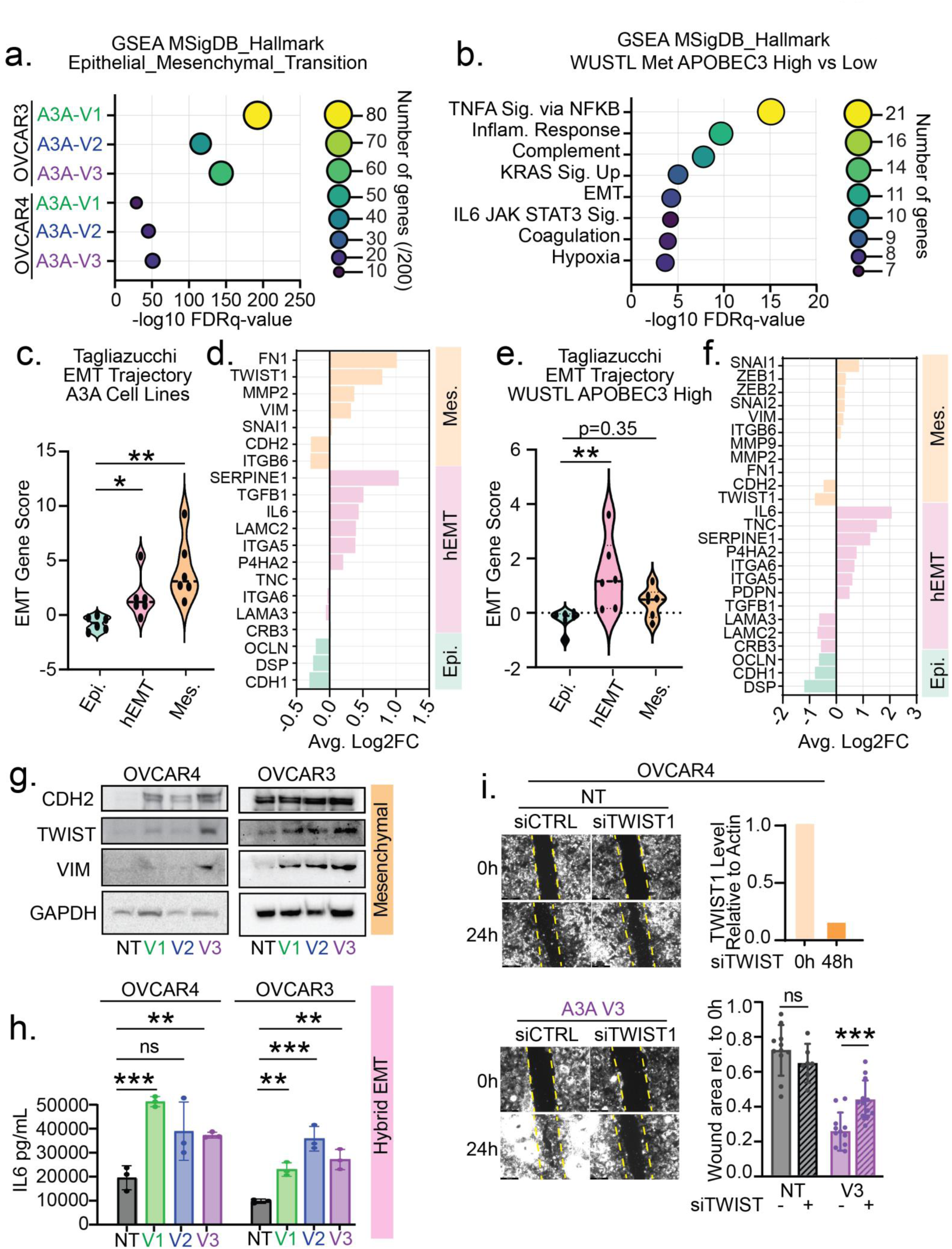
Episodic A3A alters EMT trajectory in OVCAR3 and OVCAR4 cells. **a)** RNA-seq from OVCAR4 and OVCAR3 A3A V1-3 and NT parental cell lines was performed. Gene set enrichment analysis (GSEA) of the significant differentially expressed genes (DEGs) between A3A-exposed and NT cells demonstrated that the most significantly enriched MSigDB Hallmark gene set for each version was Hallmark EMT. Significance determined by FDR q-value. Color and size of bubbles in plot indicates number of DEGs identified in the Hallmark EMT gene set (out of 200 genes total in gene set) for each A3A-exposed cell line. **b)** RNA-seq from WUSTL patient datasets was accessed and analyzed to determine DEGs from WUSTL Met APOBEC3 Low and WUSTL Met APOBEC3 High patient populations. Significant GSEA from DEGs determined by FDR q-value. Color and size of bubbles in plot indicates number of DEGs identified in each Hallmark gene set. **c)** DEGs were assessed using the Tagliazucchi EMT trajectory score to define epithelial (epi), hybrid (hEMT), and mesenchymal (mes) macro-states in each cell line. V1-3 for both cell lines are grouped together. Violin plots show EMT trajectory score for each associated profile. Values less than 0 represent under-represented gene groups. Significance determined by two-tailed t-test, *p≤0.05, **p≤0.01. **d)** Bar graph showing the DEGs found in the Tagliazucchi EMT trajectory score. **e)** The Tagliazucchi EMT trajectory was applied to DEGs from metastatic tumors from WUSTL cohort patients with APOBEC3 high v. low mutational burdens. Violin plots show EMT trajectory score for each associated profile. Values less than 0 represent under-represented gene groups. Significance determined by two-tailed t-test, **p≤0.01. **f)** Bar graph of WUSTL cohort patient DEGs from the Tagliazucchi EMT trajectory score. **g)** Immunoblot of mesenchymal markers CDH2, TWIST1, and Vimentin in OVCAR4 and OVCAR3 A3A V1-3 and NT cells. GAPDH is a loading control. **h)** ELISA for IL-6 secretion in the media of OVCAR4 and OVCAR3 A3A V1-3 and NT cell lines after 3 days in culture. Error bars are mean with SD for n=3 biological replicates. Significance was determined by two-tailed t-test, ***p≤0.001, **p≤0.01. **i)** Wound healing assay of OVCAR4 NT and OVCAR4 A3A-V3 cells depleted of TWIST1 by siRNA. Images were acquired with 4x objective, representative images are shown. Bottom right: Wound area was calculated and is plotted as 24h area relative to 0h. Significance was determined by unpaired two-tailed test, ***p≤0.001. Error bars are mean with SD for n=3 biological replicates. Top right: qPCR analysis of TWIST1 expression at 48h after knockdown, transcript levels are shown relative to actin.

EMT is a spectrum of transcriptomic and phenotypic states, the consequences of which vary with regard to progression of HGSOC(38, 51–53). We utilized a previously defined EMT gene score(38, 52) to identify how transcriptional changes resulted in epithelial, mesenchymal, or hybrid epithelial-mesenchymal (hEMT) macro-states within OVCAR3 and OVCAR4 cells exposed to episodic A3A. This analysis demonstrated that exposure to episodic A3A shifted OVCAR3 and OVCAR4 cells away from epithelial states towards hEMT and mesenchymal cell states (**Fig. 5c-d**). Using RNA sequencing from patients with HGSOC in the WUSTL cohort, we applied the EMT gene score to metastatic tumors with low or high levels of APOBEC3 mutagenesis (**Fig. 5e**). We found that high levels of APOBEC3 mutagenesis correlated with an increase in both hEMT and mesenchymal genes in metastatic HGSOC sites (**Fig. 5e-f**). We next analyzed EMT protein expression and found upregulation of mesenchymal markers (CDH2, VIM, TWIST1; **Fig. 5g**). The cytokine interleukin 6 (IL-6) is secreted by cancer cells undergoing EMT and in turn can active EMT plasticity(54–56). We found increased IL-6 secretion by in OVCAR3 and OVCAR4 cells after episodic A3A expression relative to NT controls (**Fig. 5h**), indicating the potential for EMT plasticity. Interestingly, enrichment of IL-6 signaling was also reflected in the analysis of RNA sequencing from metastatic patient tumors with high levels of APOBEC3 mutagenesis (**Fig. 5b**). These data show a correlation between APOBEC3 activity and altered EMT in both metastatic patient tumors and experimental models of HGSOC.

We hypothesized that gene and protein expression changes indicative of hEMT/mesenchymal phenotypes were responsible for the migratory phenotypes observed in A3A-exposed HGSOC cells and xenografts (**Fig. 2, 3**). We performed siRNA knockdown of the mesenchymal marker TWIST1. TWIST1 is a highly conserved transcription factor that plays a prominent role in EMT(57–65) and was upregulated in HGSOC cells exposed to episodic A3A (**Fig. 5d, 5g**). We depleted TWIST1 in OVCAR4 cells and found that migration into a wound was significantly inhibited in A3A-exposed cells whereas NT controls were unaffected (**Fig. 5i**). Taken together, we demonstrate that A3A expression alters EMT states, resulting in phenotypic changes that impact metastatic progression of HGSOC.

## DISCUSSION

Despite advances in diagnostic and therapeutic approaches, HGSOC remains the most lethal gynecologic malignancy, largely due to its propensity for metastasis and chemoresistance(2–5). Utilizing unique precursor, primary, and metastatic cancer samples from individual patients, we find a substantial contribution of APOBEC3 mutagenesis in HGSOC genomes, particularly in metastatic sites. We additionally establish a correlation between high APOBEC3 mutagenesis and decreased patient survival. Previous studies correlating elevated expression of A3B in HGSOC and patient survival reported conflicting results(9, 66). However, recent reports indicate that A3A is the more frequent contributor to APOBEC3-mediated mutations in human tumors(8, 30). We demonstrate that A3A expression in HGSOC cells promotes metastatic phenotypes and expression of EMT pathways. Notably, our murine studies reveal that A3A expression enhances distant metastatic seeding in the lungs. Thus, we define both in patient tumors and experimental models how APOBEC is associated with HGSOC progression.

The genomic landscape of HGSOC is defined by a number of recurrently mutated genes most notably *TP53* and *BRCA1/2*(43, 67, 68). However, drivers of disease progression and chemoresistance remain enigmatic. Comparison of metastatic to primary HGSOC has revealed an increase in tumor mutational burden and overall genomic instability as tumors progress(69). Here we define cytidine deamination by APOBEC3 enzymes as a mediator of mutagenesis in HGSOC, in particular in the progression from primary to metastatic tumors. We found that mutations acquired in HGSOC cells exposed to A3A recapitulated previously reported mutations in patients with HGSOC (*BRCA2*(69), *LEPR*(70), *NSD2*(71), *ITGB2*(72), *TTN*(73); **Supp. Table 1**). Interestingly, we detected a previously identified hotspot for A3A mutagenesis in the helical domain of PIK3CA (c.1624G > A; c.1633G > A)(12, 74) (**Supp. Table 1**) which is typically associated with clear cell ovarian carcinoma. Several pathogenic gene mutations attributable to A3A in our study are less frequently reported in patients with HGSOC (*TNRC6A, BMPR1B, JARID2, CUX1, PRKCG, LAMP2*; **Supp. Table 1**). Our findings of variable mutations even in isogenic models are consistent with the stochastic nature of A3A activity(7, 30) and indicate that A3A may contribute to the well-established inter-and intra-tumor heterogeneity of HGSOC(75–78).

Despite the diversity in the total number and location of mutations mediated by A3A activity across cell lines, we found that A3A activity consistently altered the phenotypes of HGSOC cells. HGSOC cells exposed to A3A and patient tumors with high levels of APOBEC3 mutagenesis exhibited consistent alterations in EMT gene expression. EMT pathways are essential in non-malignant cells during development and wound healing(37). Interestingly, a recent analysis of differentiating keratinocytes demonstrated that upregulation of A3A occurred simultaneously with genes involved in the wound healing response(46). While previous studies have identified tumor samples in which both APOBEC3 mutational signatures and EMT transcriptional profiles exist concurrently(38, 79, 80), an experimental link between A3A and EMT has not been previously reported. EMT plasticity, in which tumor cells move fluidly between epithelial and mesenchymal phenotypes, is an established mode of ovarian tumor progression and metastasis(38, 39, 47, 51, 81, 82). Importantly, HGSOC cells exhibit transcriptional profiles consistent with a combination of epithelial and mesenchymal phenotypes (hEMT) that correlate with more aggressive disease states(38, 53). Here we find that APOBEC3 activity leads to hEMT/mesenchymal transcriptional profiles in HGSOC cells and metastatic patient tumors. Among other hEMT markers, we find upregulation and secretion of IL-6 which itself is an inducer of migratory phenotypes in HGSOC(56, 83). A3A expression has been correlated with *IL6* expression in cancer cell lines and primary macrophages(84, 85). Therefore a potential model to explain the correlation between A3A and EMT is through A3A-mediated upregulation of *IL6* which promotes induction of EMT. Our study lays a foundation to study the transcriptional circuits underlying A3A-mediated EMT and provide a basis for investigating how A3A impacts EMT in other tumor types.

The molecular mechanism by which A3A activity elicits EMT transcriptional programs was not explained by recurrent genomic mutations found in HGSOC cell lines in this study. However, it is possible that transcriptional alterations are generated by deaminase-induced changes to chromatin function. A3A can act on methylated cytidines (5mC) resulting in the direct conversion of 5mC to thymidine and loss of the DNA methylation mark(7, 86–88). Global alterations in DNA-methylation patterns, which canonically repress transcription, may impact chromatin accessibility and gene expression. Interestingly, *IL6* expression was previously shown to be upregulated in HGSOC by epigenetic alterations leading to increased chromatin accessibility(83). A3A activity may similarly alter *IL6* expression through epigenetic mechanisms. Alternatively, A3A binding to promoters may impact transcriptional control. A recent study found that A3A binds the TTTC motif within interferon-stimulated response elements (IRSE) which limits IFN-stimulated gene expression(89). A similar mechanism of A3A binding to regulatory elements may influence the expression of EMT genes. It is also possible that A3A-induced DNA breaks or methylation changes may disrupt or create transcription factor binding sites. A study of the related enzyme, A3B, in breast cancer found that deaminase activity at estrogen receptor (ER) binding sites resulted in DNA breaks that, upon repair, remodeled chromatin locally to promote ER-dependent gene expression(90). A3A may induce similar chromatin alterations through damage to the genome that ultimately alters EMT gene expression. Further research into the dynamics of chromatin architecture and epigenetic modifications influenced by A3A activity is warranted.

In conclusion, our study highlights the impact of A3A activity on HGSOC progression through promotion of metastasis via EMT. By integrating genomic analyses of primary and metastatic HGSOC tumors with experimental models, we provide mechanistic insights into the role of A3A in driving HGSOC metastasis. Our findings provide a foundation for investigating A3A as a predictive or prognostic biomarker and for identifying novel therapeutic approaches aimed at mitigating the consequences of aberrant A3A activity.

## MATERIALS AND METHODS

### Cell line whole-exome sequencing

Genomic DNA was extracted using the PureLink Genomic DNA Mini Kit (Invitrogen). Library preparation and whole-exome sequencing (WES) was performed at Washington University School of Medicine Genome Technology Access Center (GTAC). gDNA (100-250ng) was fragmented using a Covaris LE220 to achieve an approximate size of 200-375bp. Libraries were constructed using the KAPA Hyper Prep Kit (KAPA Biosystems, Cat # 7962363001) on a Perkin Elmer SciCloneG3 NGS (96-well configuration) automated workstation. Individual libraries were pooled for capture at an equimolar ratio yielding up to 5µg per pool and hybridized with the xGen Exome Research Panel v2.0 reagent (IDT Technologies) capture reagents. The libraries contain custom Illumina adapters with 10 base dual unique indexes. Pooled libraries were sequenced to generate paired end reads of 151 bases using an Illumina NovaSeq X plus instrument. Base calling and demultiplexing (to create sample-specific FASTQ files) were performed with the BCL Convert utility. This processing was performed with onboard software or off-instrument using a stand-alone DRAGEN processor using the same algorithm.

#### Alignment of WES reads and mutation calling

All samples were analyzed using a DRAGEN BioIT processor running software version 4.2.4 (Illumina) in tumor-normal mode. For all OVCAR4 and OVCAR3 WES, sequencing reads were aligned to GRCh38 reference genome and outputted in the CRAM format. To determine novel mutations acquired after A3A expression, OVCAR4-A3A V1-3 and OVCAR3 V1-3 were compared to isogenic NT controls, and any shared mutations were filtered out. The oncogenicity of shared variants between at least two samples was accessed according to the ClinGene-CGC-VICC classification guidelines, via Franklin by Genoox (https://franklin.genoox.com)(50).

### Mutational signature extraction from human cancer sequencing

Human cancer data collected from the PCAWG and TCGA databases were obtained from syn11726616. *De novo* APOBEC3 and background spectra were extracted using NMF in the R package MutationalPatterns (v3.3.1)(91). Mutational signatures were deconvoluted to the COSMIC v3.2 reference signature set. Differences in mutational burden and APOBEC enrichment between samples was investigated and visualized using the R packages ggplot2 and tidyverse.

### Cell lines and culture

OVCAR4 and OVCAR3 cell lines were gifted from the laboratory of Dr. Katherine Fuh. All cell lines were cultured in DMEM + GlutaMAX supplemented with 10% tetracycline-free FBS and 1% penicillin/streptomycin. Cells were cultured at 37°C with 5% CO_2._ The creation of OVCAR4-A3A and OVCAR3-A3A cell lines was achieved through lentiviral transduction with the pSLIK-A3A lentivector with neomycin resistance^6^. Prior to murine experiments, all cell lines were transduced with an EF1α^CBR-GFP^ lentivirus to express Click Beetle Red (CBR) luciferase and GFP, generously provided by the DiPersio lab. After transduction, cells were sorted via flow cytometry to ensure that the cell population was 100% CBR-GFP positive. Cells were tested for mycoplasma at least every six months.

#### Episodic A3A induction

To achieve episodic A3A expression, 0.5 g/mL doxycycline was added to the culture media once per week, with a media change on day 3. The dox treatment regimen was repeated for 7 weeks. Three iterations of each cell line were generated by independent seeding prior to the first dox treatment and parallel culture throughout subsequent treatments (V1-3).

#### Short-intefering RNA transfection

Pooled siRNA oligonucleotides (25pmol) targeting TWIST1 (Horizon SMARTpool) were transfected into OVCAR cells using the RNAiMAX transfection reagent (Invitrogen) according to the manufacturer’s protocol. Gene depletion was confirmed by quantitative PCR.

### Quantitative PCR analysis

RNA was harvested from cell pellets using the Monarch Total RNA Miniprep Kit (New England Biolabs). cDNA was produced using the Invitrogen RNA-to-cDNA kit. Quantitative PCR was performed using PowerSyber Green PCR Master Mix (appliedbiosystems) on a QuantStudio 6 Pro (appliedbiosystems) and analyzed by Design and Analysis Software (Thermofischer).

### Immunoblotting

Cell lysates were prepared by boiling in 1X LDS (Invitrogen) for 15 minutes. Once cooled, 20% by volume beta-mercaptoethanol was added to the lysates. Samples were run on 10% bis-acrylamide gels in MOPS buffer (Invitrogen) and transferred to nitrocellulose membranes (GE) using a Bio-Rad turbo blot transfer machine. Blots were blocked in 5% milk and probed with primary antibodies (HA-BioLegend Clone HA.11, H3-Abcam, CDH2-Sigma-Aldrich, TWIST1-Santa Cruz, Vimentin-Cell Signaling Technology, GAPDH-GeneTex) overnight. Secondary antibodies for immunoblotting were obtained from Jackson ImmunoResearch (goat anti-rabbit IgG, goat anti-mouse IgG). Immunoblots were developed using ECL (Invitrogen) and analyzed on a Bio-Rad ChemiDoc MP.

### Proliferation and colony formation assays

For proliferation assays, cells were seeded in a 24-well plate and harvested and counted in triplicates on days 0, 2, 4, 6, 8 and 10 using the Countess II (Invitrogen). For colony formation assays, 2500-5000 cells were seeded in 6-well plates and allowed to grow for 14 days. Colonies were stained with a crystal violet solution and then imaged. Colonies were analyzed by ImageJ using the 3D Objects Counter with the minimum size set to 5.

### Wound healing assay

1×10^6^ cells were seeded in a 6-well plate and allowed to grow to confluence, approximately 24 hours. Media was removed from the wells and a p10 pipette tip was used to generate a cross through the well. Images were acquired at 0, 24, and 48 hours, using the cross section to ensure consistency in image acquisition. Wound closure was determined by assessing the area of the wound via a custom Cell Profiler pipeline. Wound area at 24 or 48 hours was divided by the wound area at 0 hours to determine relative wound closure.

### Spheroid invasion assay

Spheroids were created by seeding 300 cells in an ultra-low attachment round bottom 96-well plate with 1mg/mL fibronectin. Spheroids were allowed to form over 24 hours and then moved onto a flat bottom 96 well plate coated in 0.5 mg/mL Matrigel in standard culture media. Images of the spheroids were acquired at 0h, 24h, and 4d. Images were uploaded to ImageJ and the area of the spheroid was determined using the Analyze>Measure tool. The invasion area was determined by dividing the area of the spheroid at the later time point (24 hours or 4 days) to the area of the spheroid at 0 hours.

### Xenograft models

Animal protocols were compliant with Washington University School of Medicine Animal Studies Committee regulations (IACUC protocol 21-0230). Eight-week-old immunodeficient (NOD/SCID) female mice were procured from Jackson Laboratory and were housed in a sterile barrier facility prior to the start of the experiments. OVCAR cell lines were suspended in 50% Matrigel in sterile PBS and injected into the ventral left side of the peritoneal cavity. All mice were engrafted with 1×10^7^ cells in 100 L total volume. Weight, abdomen width, and tumor burden were assessed weekly.

#### Bioluminescent Imaging

Tumor growth and distribution were assessed by BLI with a Spectral Imaging AMI. Mice were injected IP with D-luciferin (150 μg/g in PBS) and imaged 10 min later with the AMI (1s exposure, binning 8, f-stop 1). Total photon flux (photons per second) was quantified using the Spectral Imaging Aura software. The region of interest was consistent throughout the experiment and encompassed the entire field of view for each mouse.

#### Tumor assessment and histopathology

At the time of death or 165 days after IP tumor injection, total tumor burden was assessed through macroscopic dissection. Distant metastatic tumor nodules to the lungs were counted. Histopathology for the lung tissue was performed by Washington University School of Medicine Anatomic and Molecular Pathology Core labs. Hematoxylin and eosin staining, WT1, and TP53 immunohistochemistry were used to determine the presence of ovarian tumor within the lung tissue. All histopathology was assessed and imaged in collaboration with an anatomic pathologist with expertise in gynecologic malignancies.

### RNA extraction and sequencing

RNA was harvested from cell pellets using the Monarch Total RNA Miniprep Kit (New England Biolabs). Library preparation and sequencing was performed at Washington University School of Medicine GTAC.

#### Library preparation

Total RNA integrity was determined using Agilent Bioanalyzer. Ribosomal RNA was removed by an RNase-H method using RiboErase kits (Roche). mRNA was reverse transcribed to yield cDNA using SuperScript III RT enzyme (Life Technologies, per manufacturer’s instructions) and random hexamers. A second strand reaction was performed to yield ds-cDNA. cDNA was blunt-ended and 3’ A-tailed followed by ligation of Illumina sequencing adapters to the ends. Ligated fragments were then amplified for 12-15 cycles using primers incorporating unique dual index tags. Fragments were sequenced on an Illumina NovaSeq using paired end reads extending 150 bases.

#### Differential gene expression analysis

RNA-seq reads were aligned to human reference genome (GRCh38.p14) with STAR version 2.5.1a. All gene counts were then imported into the R/Bioconductor package EdgeR and TMM normalization size factors were calculated to adjust for samples for differences in library size. Technical triplicates for each cell line were assessed by RNA-sequencing and the average expression for each gene identified was calculated. Differentially expressed genes (DEGs) were identified using DESeq2. DEGs were filtered using an adjusted p-value of <0.01 and fold change of >1 or <-1 for OVCAR3-A3A V1-3 and >0.5 or <-0.5 for OVCAR4-A3A V1-3. For WUSTL Met patient cohorts, DEGs were identified for gene set enrichment analysis (GSEA) using Limma to compare average gene expression between WUSTL Met APOBEC3 High and WUSTL Met APOBEC3 Low patient datasets. For EMT Trajectory Score analysis, DEGs were identified for each patient in the WUSTL Met APOBEC3 High dataset by determining fold change for each gene relative to the average expression of the gene in the WUSTL Met APOBEC3 Low patient dataset (fold change= (WUSTL Met APOBEC3 High expression-WUSTL Met APOBEC3 Low expression average)/WUSTL Met APOBEC3 Low expression average).

#### Gene set enrichment analysis

Gene set enrichment analysis (GSEA) was performed to identify MSigDB Hallmark gene sets(92). The expression value for each gene identified in Hallmark epithelial-mesenchymal-transition gene set was assessed and a heatmap was generated using Phantasus as described previously(93).

#### EMT Trajectory Score

The EMT trajectory score was derived from expression data from a curated list of genes associated with epithelial, hybrid-EMT, or mesenchymal cancer cell phenotypes, as previously described(38). The expression values for each gene attributed to the given phenotype was averaged, and for hybrid-EMT and mesenchymal gene scores the average expression value for the epithelial-associated genes was subtracted from the average expression from the corresponding average expression value for hybrid-EMT or mesenchymal genes as previously described(38).

### Enzyme-linked immunosorbent assay (ELISA)

Cells were grown for 72 hours after which culture media was collected and immediately assayed for IL-6 levels (Human IL-6 Quantikine Elisa, R&D Systems). Optical density at 540nm was determined using a SPECTROstar Omega (BMG Labtech). A standard curve was generated by using a serial dilution of the IL-6 standard in duplicate. Values were averaged, plotted, and a best-fit line was generated in Excel. The concentration of IL-6 in the culture media was determined using the linear trendline equation.

### Statistical analysis

All statistical tests were performed in R or GraphPad (Prism). Biological and/or technical triplicate tests were used to ensure robustness and reproducibility of data. p-values were generated using paired and unpaired two-tail t-tests. Error bars are mean with standard deviations.

## Data and materials availability

Variant calling from genome sequencing of patient tumors in WUSTL cohort, RNA sequencing and variant calling from genome sequencing of OVCAR cell lines, and all code used for the analysis are available at https://doi.org/10.5281/zenodo.12571514. RNA sequencing files from patient tumors in the WUSTL cohort are deposited in the NCBI GEO database under GSE218989.

## Acknowledgements

We thank all members of the Green and Bednarski labs and Drs. Katherine Weilbacher, Jason Weber, Christopher Maher, and Alessandro Vindigni for helpful discussions and input. We thank Angela Schab, Julie Ritchey and Dr. John DiPersio for sharing reagents and technical support. We thank Emily Kotnik for assistance with primary HGSOC sequencing data.

## Author Contributions

JMD, BRH, RAD, DFF, DL, LE performed experiments, analyzed and interpreted data. LS performed histopathological imaging and interpretation. TT, RD performed informatic experiments and statistical tests. JMD, AMG, KF, JJB, MM, RD conceptualized and designed experiments. JMD and AMG wrote the manuscript, with editing from all authors.

