## Supplemental Data for "APOBEC3A drives metastasis of high-grade serous ovarian cancer by altering epithelial-to-mesenchymal transition"

#### SUPPLEMENTARY FIGURES AND METHODS

Supp. Figure 1

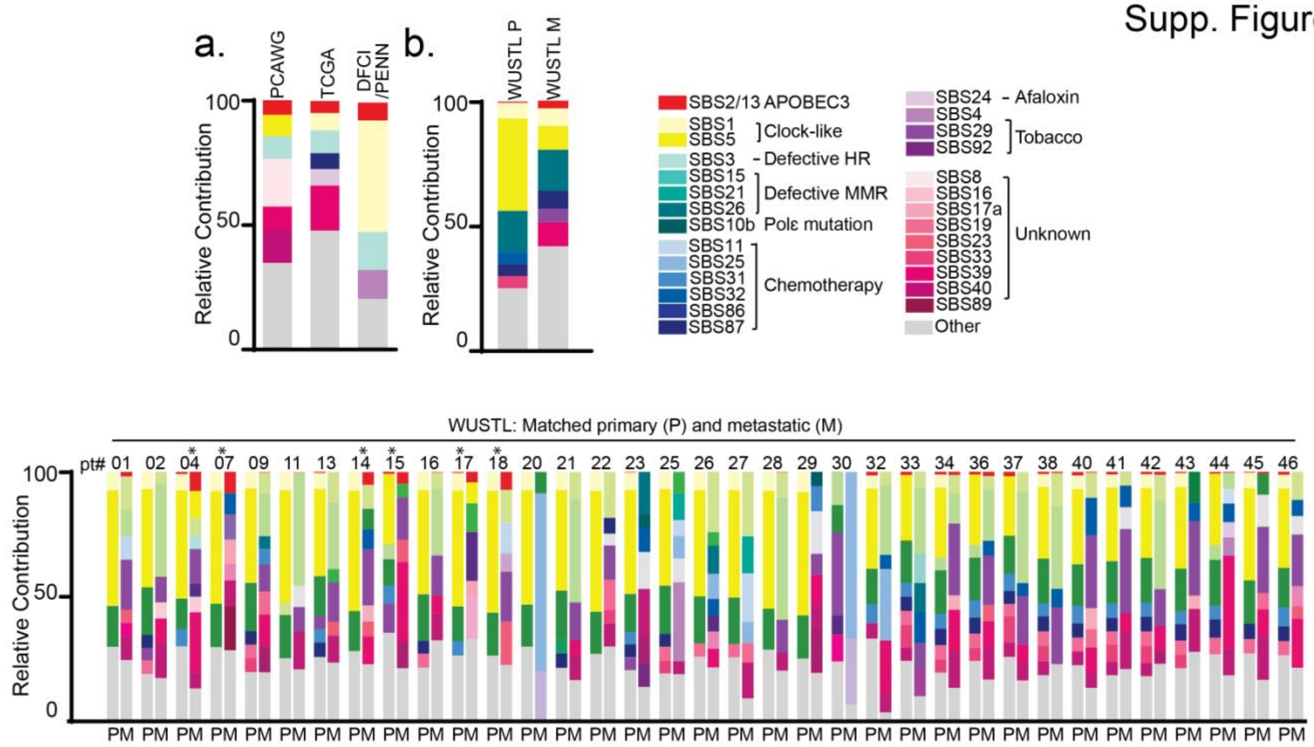


**Supp. Figure 1: SBS contribution in HGSC patient genomes. a)** Whole genome (PCAWG Ovarian Cancer) and whole exome (TCGA Ovarian Cancer, Dana Farber Cancer Institute/University of Pennsylvania) sequencing was assessed to determine the mutational processes occurring within the patient genomes. Relative contribution was determined by assessing the number of mutations identified for each SBS relative to the total mutational burden. Any mutational signature with >5% relative contribution is shown, APOBEC mutational signatures (SBS2 and SBS13) are denoted in red. Gray bars are the combination of all other signatures that were detected at <5% contribution. **b-c)** Whole exome sequencing was assessed to determine the mutational processes occurring with patient genomes from the WUSTL cohort. Any mutational signature with >5% relative contribution is shown, APOBEC mutational signatures (SBS2 and SBS13) are denoted in red. Pooled data for all 35 patients is shown in panel b. Individual primary and metastatic pairs are shown in panel c. Patients with >5% SBS2+13 relative contribution are indicated by * and are highlighted in Figure 1. Legend indicates SBS and proposed etiology for mutational signatures identified to have >5% relative contribution.

### Supp. Figure 2

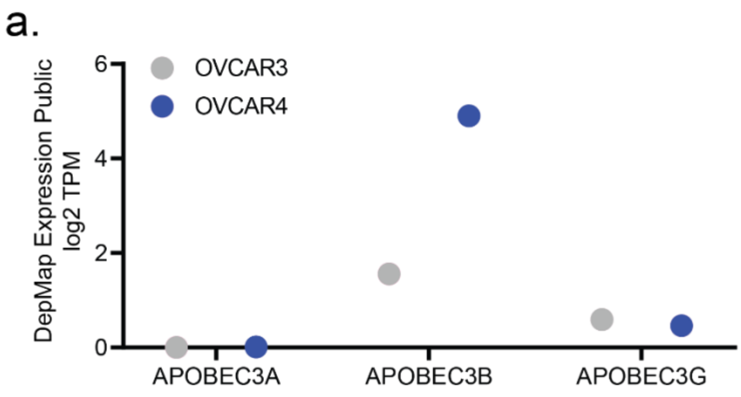


**Supp. Figure 2**: **Endogenous A3A expression is not detected in OVCAR3 and OVCAR4 cells. a)** DepMAP analysis of APOBEC3A, APOBEC3B, and APOBEC3G gene expression in OVCAR3 and OVCAR4 cell lines (Expression Public 23Q4 database). Values are log2 transcript count per million (TPM). **b)** OVCAR3 and OVCAR4 cell lines were treated with type I IFN for 24h. Cells were harvested and transcripts for APOBEC3 family members (A3A, A3B, A3G) were assessed relative to untreated controls.

### Supp. Figure 3

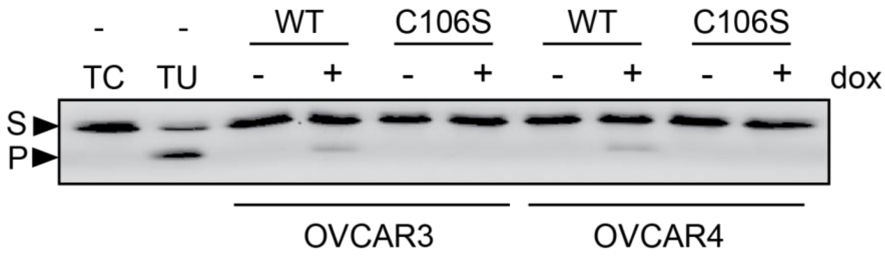


**Supp. Figure 3: A3A is active following dox induction. a)** Deaminase activity in OVCAR4 and OVCAR3 cells induced with dox to express wild-type (WT) A3A or the catalytic mutant (C106S). Lysates were incubated with a ssDNA oligonucleotide containing a single cytosine. Cytosine deamination followed by addition of uracil-DNA glycosylase (UDG) results in an abasic site; incubation with NaOH results in oligonucleotide cleavage. Substrate (S) and product (P) bands are visualized by gel electrophoresis. Oligonucleotides that contain a single cytosine (TC) or single uracil (TU), incubated in the absence of cell lysate, are used as negative and positive controls, respectively. Image is representative of n=3 biological replicates.

Supp. Figure 4

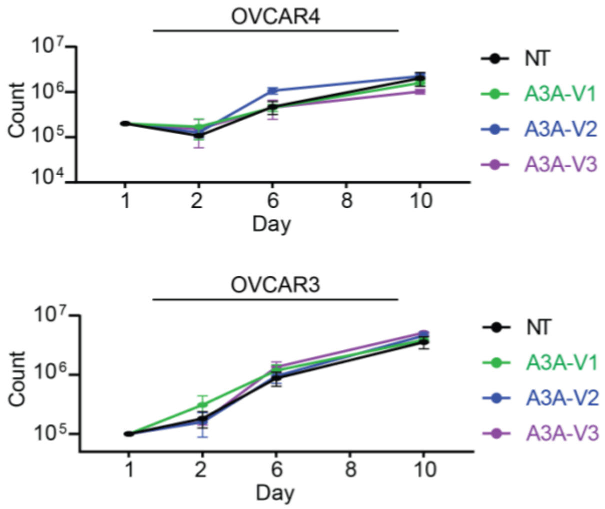


**Supp. Figure 4: A3A expression does not alter cellular proliferation in OVCAR4 and OVCAR3 cells. a)** Cells were counted on days 1, 2, 6, 8, and 10 after seeding. NT cells were cultured in parallel and counted at the same time points. Results are mean of biological triplicates. Error bars are standard deviation.

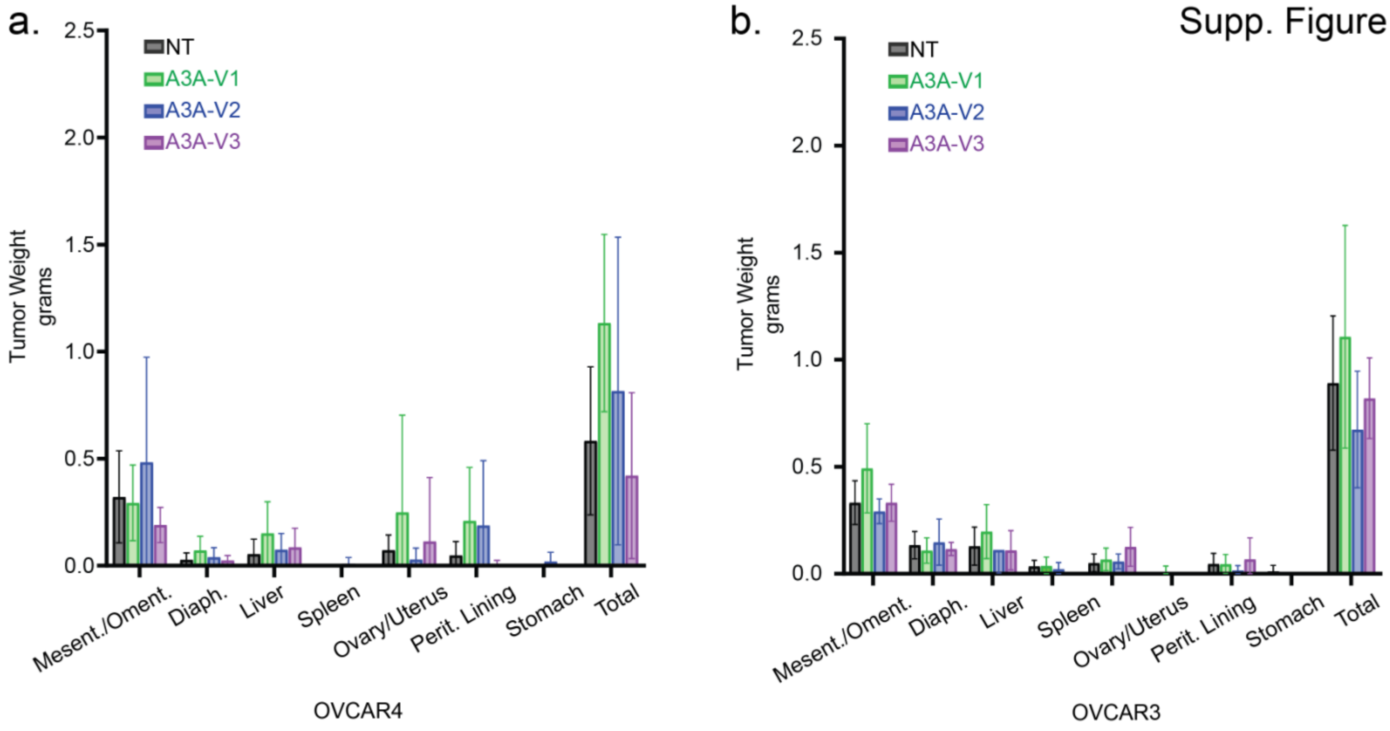


**Supp. Figure 5. Intraperitoneal tumor burden of OVCAR4 and OVCAR3 tumors. a-b)** Tumor nodules were isolated from each organ and weighed. Compiled tumor weight from each organ is shown. Error bars are mean with standard deviation. No statistically significant differences (by t-test) were detected from evaluation of A3A-exposed cells compared to NT.

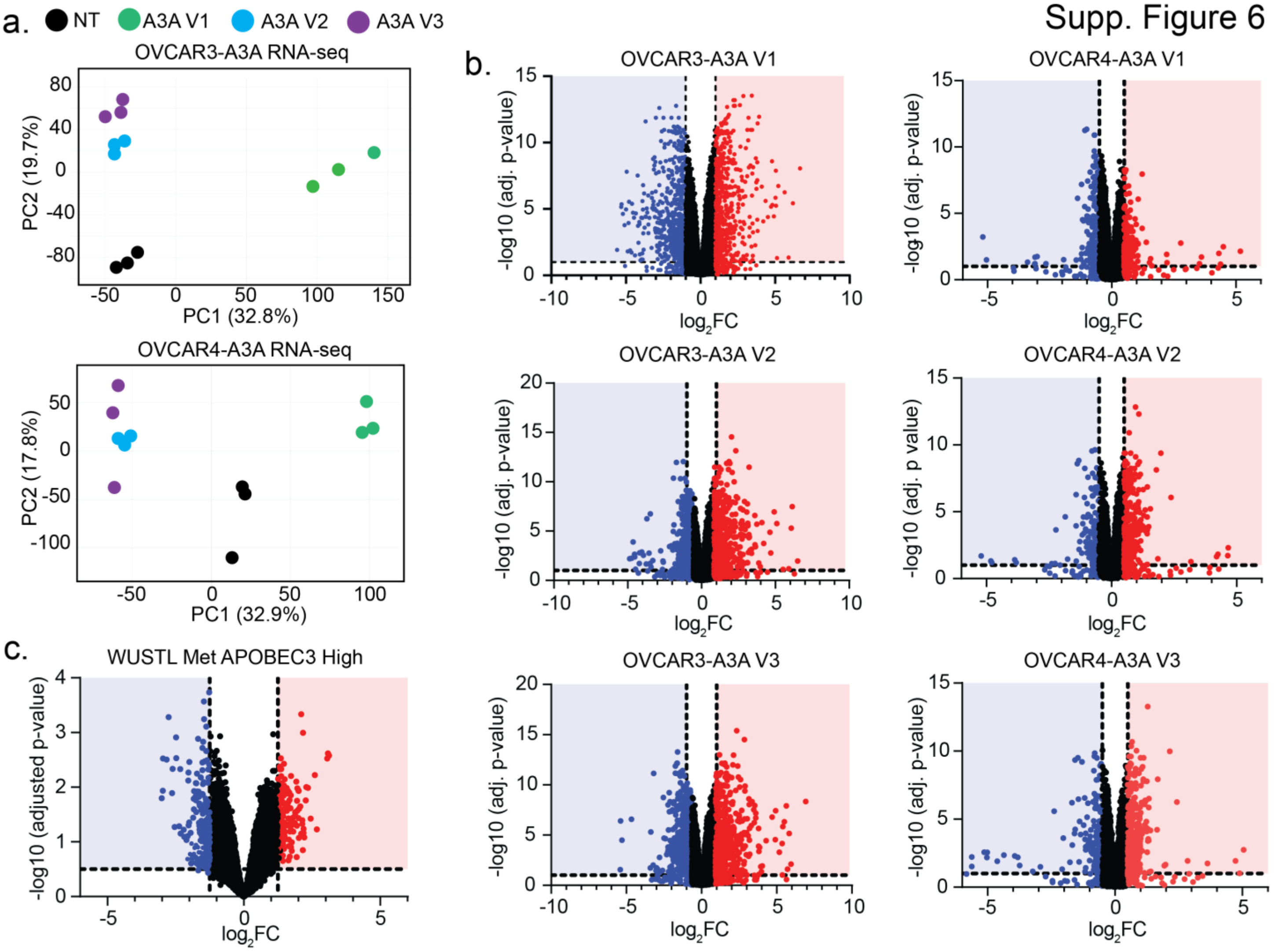


**Supp. Figure 6. Gene expression changes elicited by A3A in HGSOC cells. a)** PCA plot of gene expression in OVCAR3-A3A and OVCAR4-A3A NT and V1-3 show clustering of replicates within each sample and variance among independently-derived cell lines. **b)** Volcano plots based on RNA-sequencing data from OVCAR3-A3A and OVCAR4-A3A cells. Gene expression is plotted for each A3A-exposed version relative to NT cells. Significance threshold is set to adjusted p-value<0.01, log2 Fold Change (FC) is set to <-1 or >1 for OVCAR3-A3A and <-0.5 or >0.5 for OVCAR4-A3A. **c)** Volcano plot based on RNA-sequencing data from WUSTL cohort patients. Differential expression of genes was compared between metastatic sites with APOBEC3 High or APOBEC3 Low mutational burdens as defined in Fig. 1c. Significance threshold is set to adjusted p-value<0.01, log2 Fold Change (FC) is set to <-1.25 or >1.25.

#### SUPPLEMENTARY METHODS & REFERENCES

**DepMap APOBEC3 expression analysis.** Published expression data for OVCAR4 and OVCAR3 cell lines was first assessed using DepMap Portal for APOBEC3A, APOBEC3B, and APOBEC3G gene expression in the Expression Public23Q4 database.

**Deaminase assay**. OVCAR4-A3A and OVCAR3-A3A cells were treated with doxycycline for 24 hours then collected by centrifugation and lysed in a buffer containing 1X RIPA buffer (Cell Signaling Technology) with the addition of 1X Pierce protease inhibitor (EDTA Free, Thermo Scientific) and PMSF (GoldBio) on ice for ten minutes followed by brief sonication. Protein concentration was determined by Bradford assay. A deaminase reaction buffer containing 20mM MES and 0.1% Tween was prepared in water and the pH was adjusted to 5.9-6.1. Cell lysate was combined with reaction buffer and an oligo containing a single cytosine base (TGAGGAATGAAGTTGATT**C**AAATGTGATGAGGTGA) with a 5'-FAM fluorophore as previously described(DeWeerd and Green 2022). The negative control reaction consisted of the buffer and single-cytosine oligo, the positive control contained an identical oligo with a uracil in place of the cytosine. All samples and control reactions were incubated at 37°C for 2h, followed by the addition of 2.5 units of uracil DNA glycosylase (NEB) and subsequent at 37°C for 15m. A loading dye solution containing formamide, sodium hydroxide, and EDTA with bromophenol blue was added and reactions were boiled at 95°C for 15m. Reactions were run out on a urea-acrylamide gel in 1X TBE and the gel was imaged using the fluorescein channel on a Bio-Rad ChemiDoc MP imager.
